# Diverse motif ensembles specify non-redundant DNA binding activities of AP-1 family members in macrophages

**DOI:** 10.1101/345835

**Authors:** Gregory J. Fonseca, Jenhan Tao, Emma M. Westin, Sascha H. Duttke, Nathanael J. Spann, Tobias Strid, Zeyang Shen, Joshua D. Stender, Verena M. Link, Christopher Benner, Christopher K. Glass

**Affiliations:** University of California San Diego, School of Medicine, Department of Cellular and Molecular Medicine, La Jolla, CA 92037, USA; University of California San Diego, School of Medicine, Department of Medicine, La Jolla, CA 92037, USA; University of California San Diego, Jacobs School of Engineering, Department of Bioengineering, La Jolla, CA 92037, USA; Faculty of Biology, Division of Evolutionary Biology, Ludwig-Maximilian University of Munich, Munich, Germany

**Author notes:** these authors contributed equally to this work (Gregory J. Fonseca, Jenhan Tao).

## Abstract

Mechanisms by which members of the AP-1 family of transcription factors play both redundant and non-redundant biological roles despite recognizing the same DNA sequence remain poorly understood. To address this question, we investigated the molecular functions and genome-wide DNA binding patterns of AP-1 family members in macrophages. ChIP-sequencing showed overlapping and distinct binding profiles for each factor that were remodeled following TLR4 ligation. Development of a machine learning approach that jointly weighs hundreds of DNA recognition elements yielded dozens of motifs predicted to drive factor-specific binding profiles. Machine learning-based predictions were confirmed by analysis of the effects of mutations in genetically diverse mice and by loss of function experiments. These findings provide evidence that non-redundant genomic locations of different AP-1 family members in macrophages largely result from collaborative interactions with diverse, locus-specific ensembles of transcription factors and suggest a general mechanism for encoding functional specificities of their common recognition motif.

## Introduction

Gene expression is controlled by sequence-specific transcription factors (TFs) which bind to promoters and distal enhancer elements^1-3^. Genome wide studies of regulatory regions in diverse cell types suggest the existence of hundreds of thousands of enhancer sites within mammalian genomes. Each cell type selects a unique combination of ~20,000 such sites that play essential roles in determining that cell’s identity and functional potential^4-7^. Selection and activation of cell-specific enhancers and promoters is achieved through combinatorial actions of the available sequence-specific TFs^8-14^.

TFs are organized into families according to conserved protein domains including their DNA binding domains (DBD)^15^. Each family may contain dozens of members which bind to similar or identical DNA sequences^16,17^. An example is provided by the AP-1 family, which is composed of 15 monomers subdivided into five subfamilies based on amino acid sequence similarity: Jun (Jun, JunB, JunD), Fos (Fos, FosL1, FosL2, FosB), BATF (BATF, BATF2, BATF3), ATF (ATF2, ATF3, ATF4, ATF7) and Jdp2^18-22^. AP-1 binds DNA as an obligate dimer through a conserved bZIP domain. All possible dimer combinations can form with the exception of dimers within the Fos subfamily^23^. The DBD of each monomer of the AP-1 dimer recognizes half of a palindromic DNA motif separated by one or two bases (TCASTGA and TCASSTGA)^16,17,24-26^. Previous work has shown that dimers formed from Jun and Fos subfamily members bind the same motif^16^. Given a conserved DBD, and the ability to form heterodimers, it naturally follows that different AP-1 dimers share regulatory activities. However, co-expressed family members can play distinct roles. For example, Jun and Fos are co-expressed during hematopoiesis, but knockout of Jun results in an increase in hematopoiesis whereas knockout of Fos has the opposite effect^27-30^. The basis for non-redundant activities of different AP-1 dimers and heterodimers remains poorly understood.

Specific AP-1 factors have been shown to form ternary complexes with other TFs such as IRF, NFAT and Ets proteins, resulting in binding to composite recognition elements with fixed spacing^31-33^. However, recent studies examining the effects of natural genetic variation suggested that perturbations in the DNA binding of Jun in bone marrow derived macrophages are associated with mutations in the motifs of dozens of TFs that occurred with variable spacing^34^. These observations raise the general question of whether local ensembles of TFs could be determinants of differential binding and function of specific AP-1 family members. To explore this possibility, we examined the genome-wide functions and DNA binding patterns of co-expressed AP-1 family members in resting and activated mouse macrophages. In parallel, we developed a machine learning model, called a Transcription Factor Binding Analysis (TBA), that integrates the affinities of hundreds of TF motifs and learns to recognize motifs associated with the binding of each AP-1 monomer genome-wide. By interrogating our model, we identified DNA binding motifs of candidate collaborating TFs that influence specific binding patterns for each AP-1 monomer that could not be identified with conventional motif analysis. We confirmed these predictions functionally by leveraging the natural genetic variation between C57BL/6J and BALB/cJ mice, and observing the effects of single nucleotide polymorphisms (SNPs) and short insertions or deletions (InDels) on AP-1 binding. Finally, we confirm the model’s prediction of PPARγ binding being specifically associated with the selection of a single family member, Jun, using PPARγ-deficient macrophages.

## Results

### AP-1 family members have distinct regulatory functions in macrophages

AP-1 family members are ubiquitously expressed with each cell type selecting a subset of family members (monomers), which make up the AP-1 dimer. Each family member shares a conserved DNA binding and dimerization domain but are dissimilar outside of the basic leucine zipper (bZIP domain, Fig 1A). RNA-seq performed on thioglycolate-elicited macrophages (TGEMs) revealed ATF3, Jun, and JunD as the most expressed AP-1 family members under basal conditions (Veh, Fig. 1A, right, Supplementary Fig. 1A). Following activation of TGEMs with Kdo2 lipid A (KLA), a specific agonist of TLR4^35^, there is a marked increase in Fos, Jun and JunB expression, consistent with AP-1 family members having context-specific roles (Fig. 1A).

**Figure 1.**
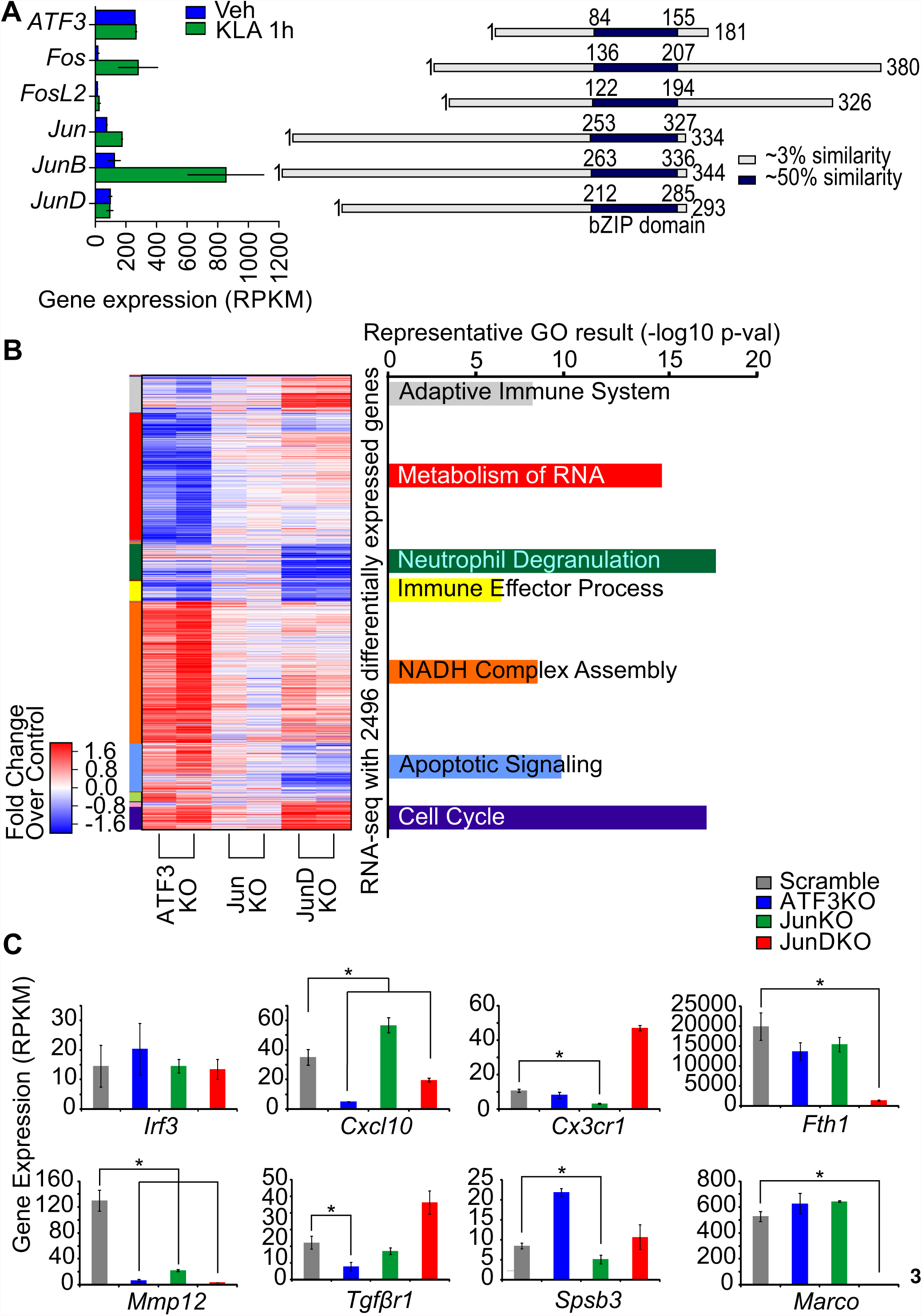
AP-1 proteins have overlapping and distinct transcriptional functions in macrophages. **A**. Protein alignment of monomers (right) and mRNA expression of monomers in TGEMs before and after 1-hour KLA treatment (left). **B.** Hierarchical clustering of genes that are differentially expressed in iBMDMs subjected to CRISPR mediated knockdown of the indicated AP-1 monomer with respect to scramble control. Expression values are given as the fold change with respect to scramble; values are Z-score normalized across each row. Representative functional annotations for each gene cluster are calculated using Metascape and the enrichment of each term is quantified as the negative log transform of the p-value. **C.** Expression of a subset of genes in AP-1 protein knockouts. * indicates FDR < 0.05.

To examine the regulatory function of individual family members, knockout cell lines for ATF3, Jun and JunD were produced using CRISPR/Cas9-mediated mutagenesis in immortalized bone marrow-derived macrophages (iBMDMs). Knockout efficiency was confirmed by western blotting (Supplementary Fig. 1B). RNA-seq analysis identified 2496 genes differentially expressed when comparing the knockout to control cells (FDR<0.05, fold change >2, RPKM≥16 Fig. 1B, Supplementary Fig. 1C). Clustering of differentially expressed genes revealed distinct clusters that were affected in individual knockout cell lines, demonstrating that each family member has distinct as well as redundant activity. The Jun knockout had a more modest effect on gene expression than the ATF3 and JunD knockout (125, 651, and 1564 differentially expressed genes respectively), suggesting that Jun may have more redundant activity (Fig. 1B and S1B). Each of the gene clusters was enriched for Gene Ontology terms for differing biological functions, including cell cycle, immune effector process and NADPH complex assembly (Fig. 1B). Examples of affected genes are shown in Figure 1C. *Mmp12* is affected by knockdown of all three factors, whereas *Marco* and *Fth1* exhibit minimal changes in expression in ATF3 and Jun KO, but decreased expression in the JunD KO iBMDMs.

### AP-1 family members can target distinct loci in addition to overlapping loci

Given the distinct roles of individual family members in regulating macrophage transcription, we used chromatin immunoprecipitation followed by deep sequencing (ChIP-seq) to map the binding of each family member in resting TGEMs treated with vehicle (Veh) or KLA for one hour (activated TGEMs). These experiments detected a substantial number of binding sites (n > 10000, IDR < 0.05) for family members with the highest mRNA expression (Supplementary Fig. 1A, Supplementary Fig. 2A). ATF3, Jun, and JunD binding sites were detected in both Veh and KLA treatment whereas Fos, Fosl2 and JunB bind predominantly after KLA treatment (Supplementary Fig. 2A). Hierarchical clustering of all 50664 AP-1 binding sites (Fig. 2A) found in either Veh or KLA treated TGEMs according to the relative binding strength of the family members (normalized to a maximum of 1 at each locus) yielded distinct subclusters that highlight the specific binding patterns of AP-1 family members as well as the reorganization of AP-1 cistromes in KLA treated macrophages (Fig. 2A). Representative regions that show distinct binding patterns of AP-1 family members are shown (Fig. 2B, Supplementary Fig. 2B).

**Figure 2.**
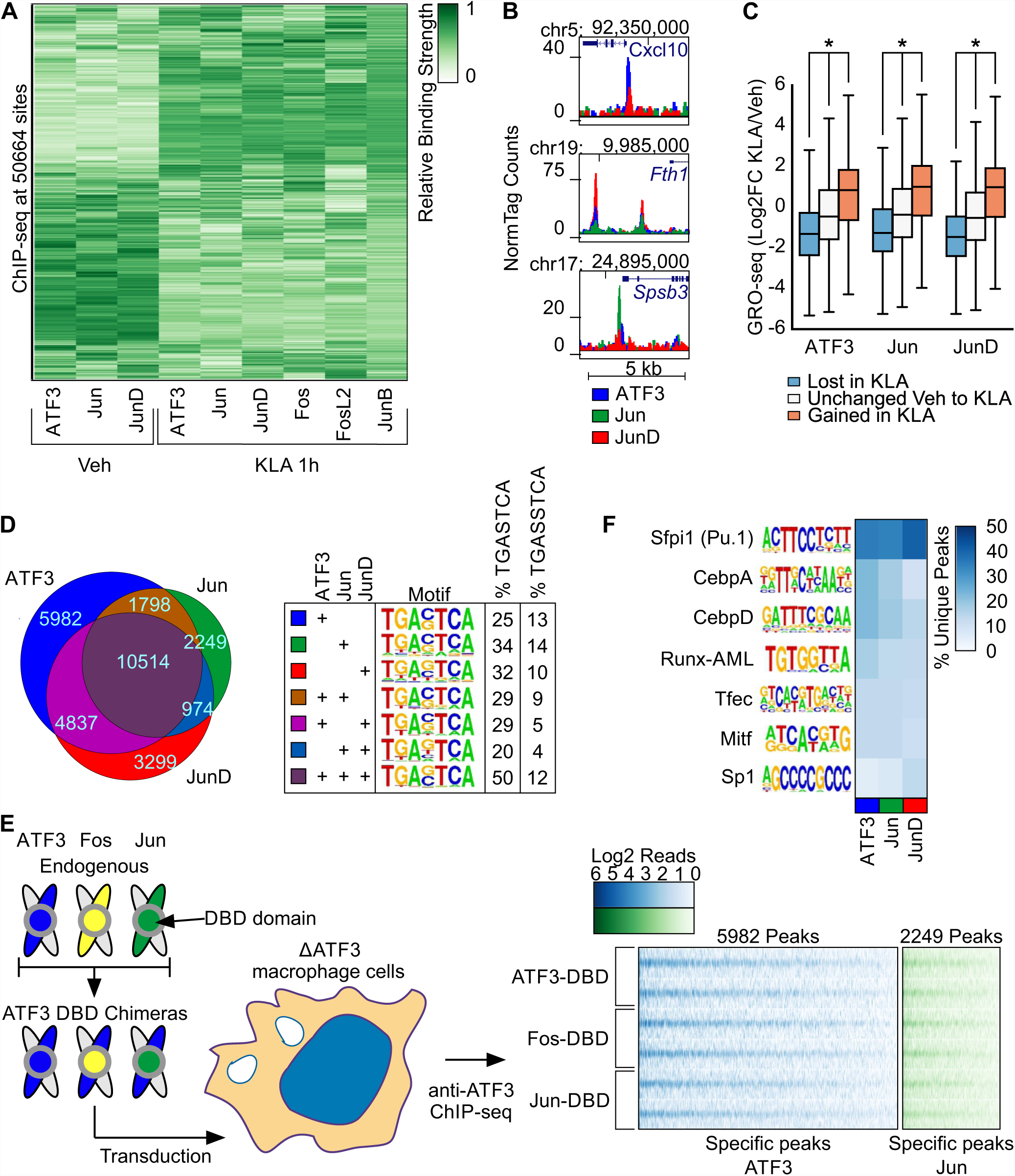
AP-1 monomers bind at unique loci that cannot be explained by differences in the DNA binding domain. **A.** Hierarchical clustering of the relative strength of binding of each monomer at all AP-1 binding sites in Vehicle and 1-hour KLA treatment conditions. **B.** Representative browser shots of ChIP-seq peaks for Veh specific monomers ATF3, Jun and JunD. **C.** GRO-seq at sites where ATF3, Jun and JunD was lost, gained or unchanged after one hour KLA treatment. **D.** Venn diagram of ATF3, Jun and JunD peaks in Vehicle (left) and table indicating the de novo AP-1 motifs found in each subset of peaks and the percent of peaks in each subset that contain one of the two AP-1 motif variants (right). **E.** Binding strength comparison of ATF3 chimeras. The ATF3 DNA binding domain (blue) is replaced the DNA binding domains of Fos (yellow) or Jun (Green) and then transduced into ATF3-deficient iBMDM cells with a lentivirus vector (left). The binding of each chimera is shown as a heatmap of ChIP-seq tags centered on ATF3 chimera binding sites (replicates indicated in separate rows) that were found to be specific for ATF3 (blue) or Jun binding in TGEMs (Green). **F.** Heatmap showing the percent of unique binding sites for each monomer that contain a de novo motif calculated from each set of unique peaks.

The gain and loss of binding sites of ATF3, Jun and JunD after KLA treatment provided an opportunity to correlate changes in their DNA occupancy with local changes in enhancer activity. Changes in the expression of enhancer-associated RNAs (eRNAs) are highly correlated with changes in enhancer function and nearby gene expression^11^. To detect eRNAs, we performed Genome Run-On Sequencing (GRO-seq) in TGEMs, which provides a quantitative measure of nascent RNA^36^. We examined GRO-seq signal at ATF3, Jun and JunD binding sites exhibiting gain, loss or no change in binding after KLA treatment. In each case, AP-1 occupancy was associated with greater GRO-seq signal (Fig. 2C). These findings suggest that ATF3, Jun and JunD primarily function as transcriptional activators.

### Family member specific binding sites are associated with the same AP-1 motif

While 10514 of the binding sites of ATF3, Jun and JunD in the vehicle condition are shared by all three factors, a greater number of binding sites (11530) are not (Fig. 2D). To ensure that the unique sites were not technical artifacts, we ranked the peaks of each family member according to the number of ChIP-seq tags detected and then calculated the percent of peaks that were unique after filtering away binding sites that fell below a given percentile threshold. We found that unique peaks were present even at higher thresholds, supporting our observation that AP-1 family members can bind to distinct loci (Supplementary Fig. 2C).

Using de novo motif enrichment analysis, we observed that the binding motif for each combination of monomers was nearly identical (Fig. 2D). To investigate whether family members preferred either variant of the AP-1 motif, we calculated the percent of peaks bound by each combination of monomers that had the TRE variant of the AP-1 motif (TGASTCA) and the CRE variant of the motif (TGASSTCA)^16,37^. Consistent with previous studies, we found both variants of the AP-1 motif at regions bound by each combination of monomers, but there was a preference for the TRE motif (Fig. 2D)^16^. These results suggest that differences in the AP-1 DBD cannot explain the majority of family member specific binding.

To test the prediction that differences in the AP-1 DBD do not explain binding patterns, we created ATF3 chimeras by replacing the DBD of ATF3 with that of Fos and Jun (Fig. 2F, S2D). The DBDs of these three factors are highly conserved, with identity at 8 and charge conservation at 3 of 11 amino acids directly involved in DNA interaction (Supplementary Fig. 2D)^24^. We transduced expression vectors for ATF3 chimeras with either an ATF3, Fos or Jun DBD into ATF3 KO iBMDMs and then measured the genome-wide binding patterns of each chimera by performing ChIP-seq using an antibody specific for ATF3 (Fig. 2F, S2D). Globally, we observed that the chimeras had stronger binding at ATF3 specific sites in comparison to Jun specific sites and that each chimera exhibited similar binding across all loci visualized as normalized tag counts in a heatmap (Fig. 2E). Representative browser shots showing similar binding between chimeras are shown at *Cxcl10* and *Spsb1* which are loci specifically bound by ATF3 and Jun respectively (Supplementary Fig. 2E).

Given that the family members all recognized a common DNA binding motif, we hypothesized that differential interactions with locally bound factors mediated by non-conserved protein contact surfaces may explain unique monomer binding sites. We calculated de novo motifs enriched at the unique peaks for ATF3, Jun, and JunD individually, and then calculated the percent of each family member’s specific binding sites that contained a match to each de novo motif. We identified motifs for key TFs in macrophages^10,34^ such as PU.1, CEBP, and Runx (Fig. 2F). Composite motifs for AP-1 and IRF or NFAT occurred at similar frequencies at the unique peaks for each family member (~5% and ~3% of peaks respectively). However, we found no significant differences in the relative enrichment of motifs associated with ATF3, Jun, and JunD specific peaks that would explain their specific binding profiles (Fig. 2F).

### A machine learning model that relates combinations of motifs to transcription factor binding

Given the robustness of the family member specific peaks (Supplementary Fig. 2C), we considered additional biological mechanisms that might be leveraged for detection of motifs differentially associated with each family member. Current methods for calculating enriched motifs analyze each motif individually despite data demonstrating that TFs bind cooperatively in groups^1,31^. Additionally, collaborative binding by TFs allows for partners to bind to more degenerate motifs, which are ignored in de novo motif analysis^10^. We incorporated these concepts into a machine learning model that relates the presence of multiple TF motifs, which may be degenerate, to the binding of a TF. Machine learning models are often considered difficult to interpret due to their complexity. In building our model, we emphasized simplicity and as a consequence, interpretability.

Figure 3A summarizes our model, TBA (Transcription factor Binding Analysis). TBA takes the binding sites of a TF as input and selects a set of GC-matched background loci. For each binding site and background locus, TBA calculates the best match to hundreds of DNA binding motifs, drawn from the JASPAR library, and quantifies the quality of the match as the motif score (aka log likelihood ratio score). To allow for degenerate motifs, all motif matches scoring over zero are considered. The motif scores are then used to train the TBA model to distinguish TF binding sites from background loci. TBA scores the probability of observing binding at a sequence by computing a weighted sum over all the motif scores for that sequence. The weight for each motif is learned by iteratively modifying the weights until the model’s ability to differentiate binding sites from background loci no longer improves. The final motif weight measures whether the presence of a motif is correlated with TF binding. The significance of a given motif can be assigned by comparing the predictive performance of a trained TBA model and a perturbed model that cannot recognize that one motif with the likelihood ratio test.

**Figure 3.**
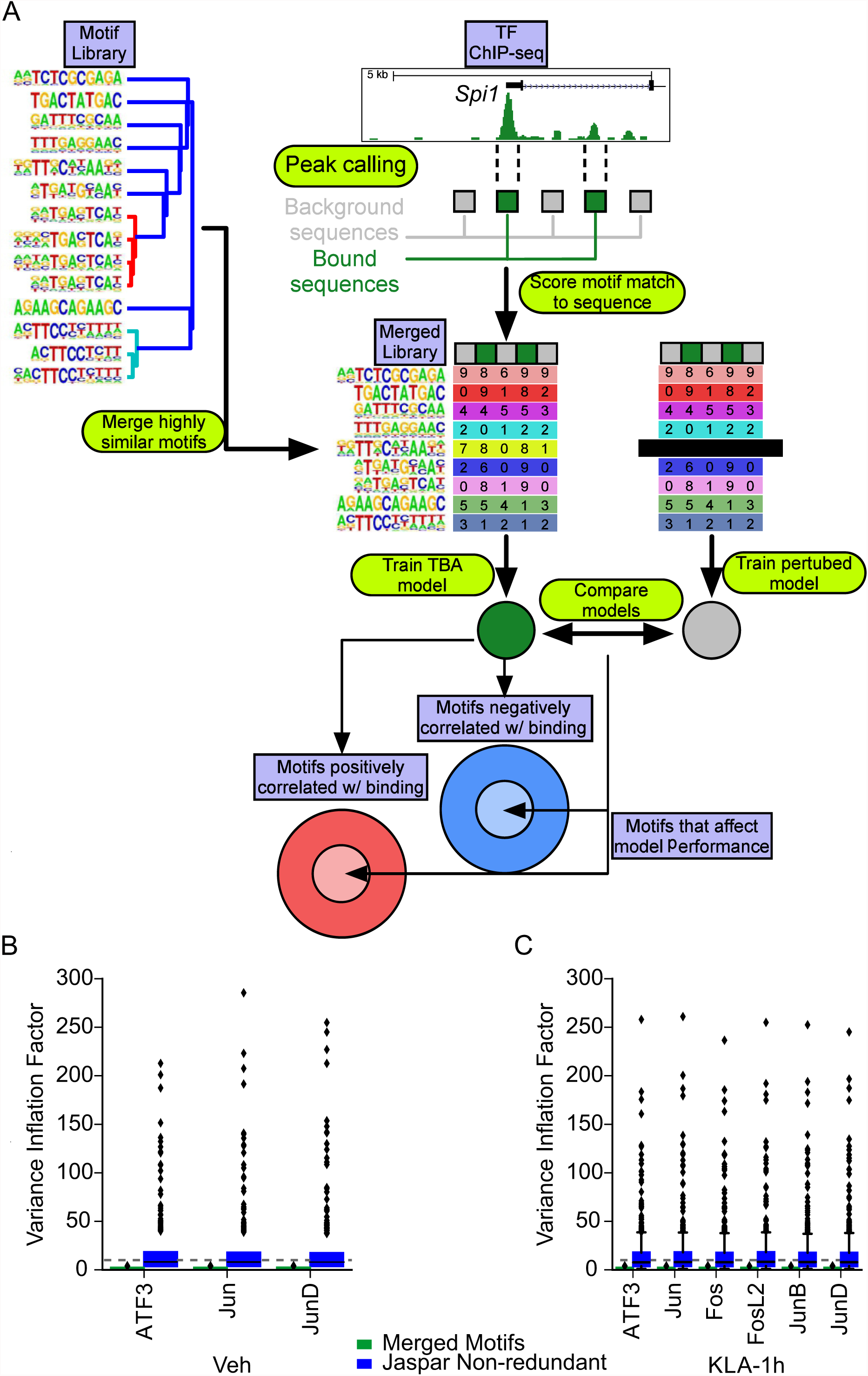
TBA, a Transcription factor Binding Analysis. **A.** Schematic workflow of TBA. Binding sites for a transcription factor (green boxes) are mixed with random GC-matched background sequences (grey boxes). Motifs from the JASPAR library are merged to create a non-redundant motif library. Motif scores are calculated for all sequences at all binding sites and GC-matched background and then used to train a TBA model. Model weights from the trained model indicate whether a motif is positively or negatively correlated with the occupancy of a transcription factor. The performance of the full model and a perturbed model with one motif removed are compared to identify motifs that are important to the model. The intersection of important motifs that affect model performance and the model weights learned by the classifier can be used to infer the binding partners of a transcription factor. **B-C**. Distribution of Variance Inflation Factor for each motif in the TBA merged motif library and JASPAR motif library for experiments performed in **(B)** Vehicle and **(C)** KLA treated TGEMs.

Machine learning models, including TBA, can be confounded by collinearity, which in our case corresponds to the presence of motifs that are highly similar or redundant^38^. Collinearity can cause inaccurate weight and significance to be assigned to motifs. To assess the extent of collinearity, we calculated the Variance Inflation Factor (VIF)^38^ for the scores of each motif in the JASPAR library at AP-1 binding sites. A VIF above 10 would indicate problematic collinearity and that the scores for a motif are highly correlated with the scores of another motif. We found that a substantial number of motifs were collinear with at least one other motif (VIF > 10) (Fig. 3B, 3C). To address the presence of redundant motifs we clustered the JASPAR library, identifying groups of motifs that are highly similar (Supplementary Fig. 3, colored clades), and merged these motifs together (Pearson Correlation > 0.9, Supplementary Fig. 3, Fig. 3), resulting in a condensed library of 196 motifs formed from 519 JASPAR motifs. Multiple collinearity was substantially reduced in our condensed library (VIF < 10, Fig. 3B, 3C).

### TBA identifies combinations of binding motifs that coordinate AP-1 recruitment

To identify motifs associated with specific AP-1 family members, we trained TBA models for each monomer in resting TGEMs, and probed for differences in the identified motifs. Ranking each motif according to the mean p-value, we found that all family members shared a core set of highly significant motifs both positively and negatively correlated with binding (Fig. 4A, i and ii, respectively). The motifs exhibiting strong positive correlation included the AP-1 motif as well as motifs of macrophage collaborative binding partners for AP-1, such as PU.1 and CEBP^10,11,34^. To determine a significance threshold for more moderately ranked motifs, we compared significance values calculated by TBA models trained on replicate ChIP-seq experiments. We determined that motifs with a mean p-value<10e-2.5 tended to have similar significance values (absolute likelihood ratio ~1, Supplementary Fig. 4A). The motif weights that exceeded this threshold were highly correlated between replicate experiments (Supplementary Fig. 4C). Outside of the core group of motifs shared by all monomers, we observed ~50 motifs with differential affinities (likelihood ratio > 100 between at least 2 monomers) for each monomer as defined by TBA (Fig. 4A, center panel, shaded regions). Differential motifs positively correlated with binding (Fig. 4A left heatmap in red) included motifs unique to a monomer such as the PPAR half site with Jun. The full PPARγ motif was negatively correlated with both ATF3 and JunD, suggesting that PPARγ positively influences the binding of Jun to a greater extent than the other AP-1 monomers (Fig. 4A right heatmap in blue). These results suggest that AP-1 monomers have distinct sets of collaborating TFs that affect their binding patterns.

**Figure 4.**
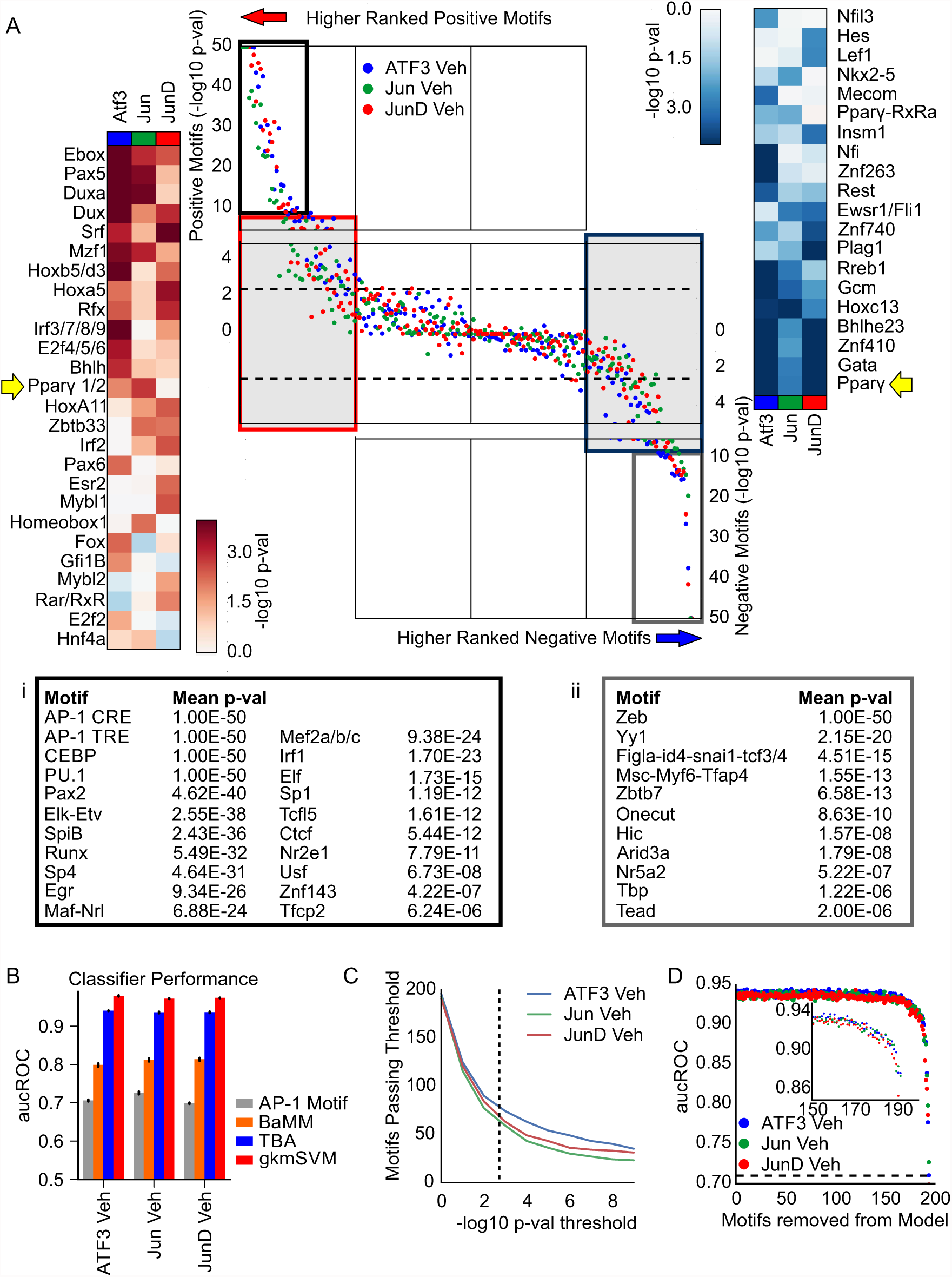
TBA identifies motifs predicted to specify differential AP-1 monomer- binding in resting TGEMs. **A.** DNA motifs rank order based on the significance of the motif according to the likelihood ratio test. The black box represents the most significant motifs positively correlated with binding for all AP-1 monomers and are listed in (i) and the most significant motifs negatively correlated with binding for all AP-1 monomers are shown in the grey box and are listed in (ii). The significance of motifs positively correlated with binding that show a 100-fold likelihood difference between two monomers are shown on the left heatmap (red); the right heatmap (blue) gives the significance of corresponding motifs negatively correlated with binding. **B.** Comparison of the performance of TBA against the AP-1 motif score alone, Bayesian Markov Model (BaMM) motif score, and gapped k-mer SVM as measured by the area under the Receiver Operating Characteristic curve (aucROC). Error bars indicate the standard deviation of aucROC across 5 cross validation sets. **C.** Number of motifs that pass an in-silico mutagenesis test for significance (the likelihood ratio test comparing the performance of a full model that uses all the motifs and a mutated model with one motif removed) at various p-value thresholds. **D.** Predictive performance of TBA when predicting ATF3, Jun and JunD binding as motifs are iteratively removed starting from the least important motif based on the weights calculated by TBA. Inset shows performance values beginning at 150 motifs removed where predictive performance begins to drop.

### Evaluation of collaborating TF motifs that coordinate AP-1 binding

To assess whether the additional motifs identified by TBA are useful for identifying AP-1 sites, we compared TBA’s ability to predict the binding of each monomer to several other sequence based approaches. Predicting TF binding using just the AP-1 TRE motif score had the worst performance as measured by the area under the receiver operating characteristic curve (aucROC (Fig. 4B). Bayesian Markov Model motifs (BaMM)^39^, which assesses dependencies between the positions within the binding motif, improved upon the simple AP-1 motif score by ~15% (Fig. 4A). The TBA model and the gkm-SVM model achieved even higher performance, demonstrating that additional sequences outside of a TF’s motif may contribute to binding site selection (Fig. 4B). The performance of gkm-SVM exceeded that of TBA (by ~3%). However, gkm-SVM cannot retrieve motifs beyond the binding motif of a single TF^40^ while TBA identified over 50 motifs that passed a significance threshold of p<10e-2.5 (Fig. 4C). To examine the impact of statistically significant (p<10e-2.5) but moderately ranked motifs, we calculated TBA’s performance while iteratively removing motifs from the model (starting with the least significant motif) (Fig. 4D). The performance of the model started declining when the motifs from the top 50 were removed, demonstrating that the local sequence environment outside of the AP-1 motif affects AP-1 binding (Fig. 4D, inset).

### Cell type specific binding preferences of JunD

To further test the hypothesis that distinct sets of collaborating TFs can affect AP-1 binding, we examined JunD binding in a panel of cell lines. Each cell type expresses a distinct repertoire of TFs that are available as binding partners for JunD. We trained TBA models for ChIP-seq of JunD in each cell line and then extracted the 20 most significant motifs from each model. Motifs which are bound by TFs known to be important for particular cell lines were found to be correlated with JunD binding. For example, the Gata motif was positively correlated with JunD binding in K562 cells, an erythroid lineage erythroleukemia, while Pou motifs (e.g. OCT4) were important in h1-hESCs (Supplementary Fig. 4C)^41^. Differences in the motifs identified by TBA for each cell line corresponded to large differences in the loci bound by JunD (Supplementary Fig. 4D).

### KLA treatment changes the collaborating TFs available to AP-1 and remodels the AP-1 cistrome

Given that AP-1 binds collaboratively with other TFs, the selection of binding sites for each monomer will depend on the available of collaborating partners. To study effects of changes in collaborating TF availability, we examined AP-1 binding before and after KLA treatment. Treatment of TGEMs with KLA resulted in 178 mRNAs increasing 2-fold (FDR<0.05) or greater (Fig. 5A). A total of 29 genes encoding TFs with known binding motifs (20 upregulated and 9 downregulated) had a significant change in expression (FDR < 0.05) including AP-1 monomers Fos, Fra2 and JunB (Supplementary Fig. 5A, blue points). In addition, TLR4 activation by KLA results in the activation of several latent transcription factors, including NFκB and interferon regulatory factors (IRFs). Correspondingly, AP-1 monomers showed changes in their global binding patterns with Fos and JunB displaying drastic upregulation in binding sites (Supplementary Fig. 2A, Fig. 5A).

**Figure 5.**
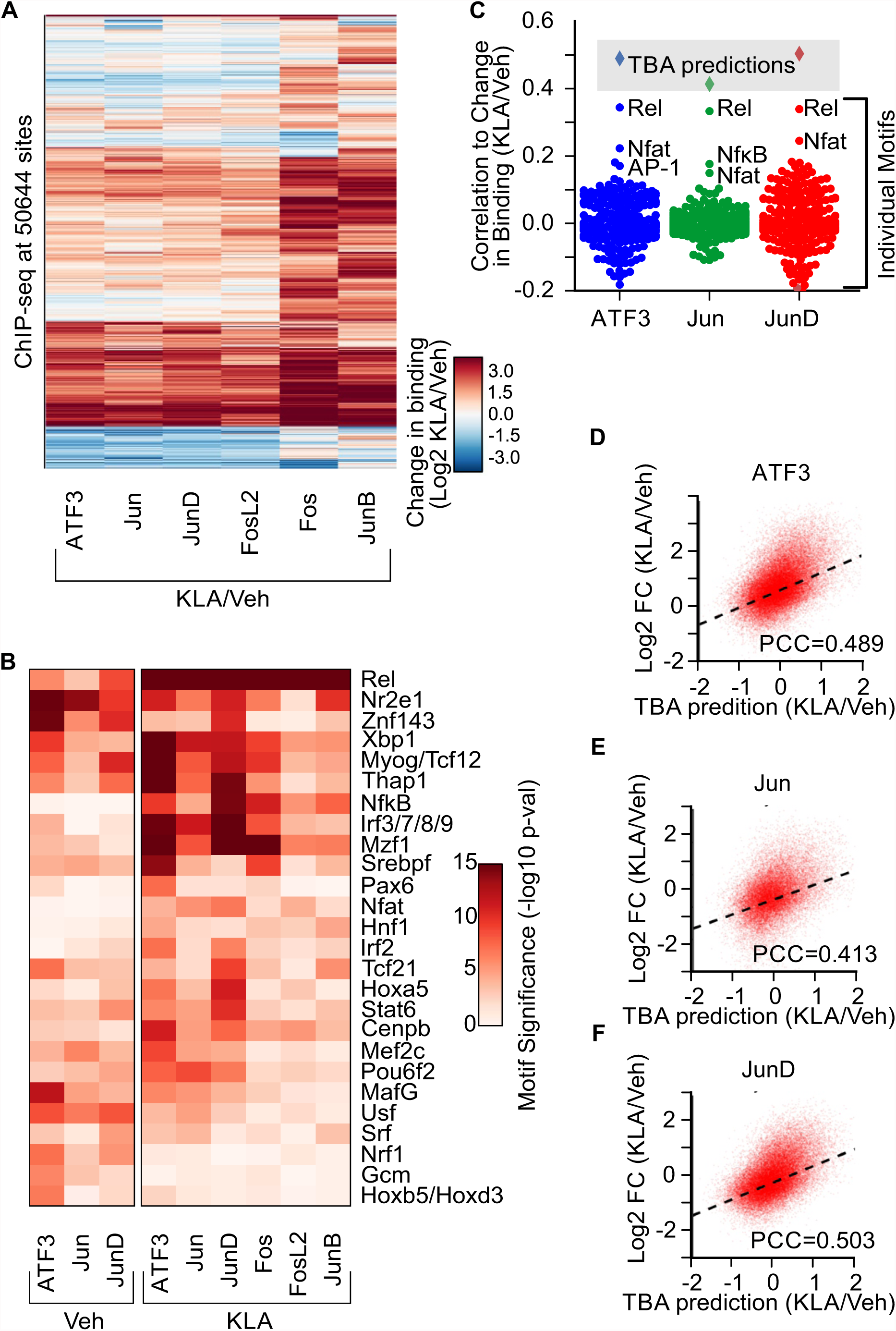
AP-1 binding is context-dependent and affected by the availability of binding partners. **A.** Heatmap of the change in binding of AP-1 monomers after 1-hour KLA treatment quantified as the Log2 ratio of KLA binding to Vehicle binding. **B.** Heatmap showing the TBA assigned significance of DNA motifs that had a 10e4 absolute likelihood ratio between the KLA and Vehicle value for each monomer. **C.** Pearson correlation of individual motif scores and TBA predictions with the change in binding after one hour KLA treatment. **D. E. F.** TBA predicted change in ATF3, Jun, and JunD binding after KLA-1h treatment versus actual change in binding. PCC indicates the Pearson Correlation coefficient of TBA predictions to the log2 fold change in binding of each monomer after one hour KLA treatment.

To examine motifs associated with AP-1 binding after KLA treatment, we trained TBA models for each monomer in KLA treated TGEMs. Again, we observed that all AP-1 monomers shared a common group of highly significant motifs positively correlated with binding, including AP-1, CEBP, PU.1, REL, and Egr, and negatively correlated with binding, such as the Zeb1 motif (Supplementary Fig. 5B, Supplementary Table 1, Supplementary Table 2). Many of the moderately ranked motifs showed large differences in significance between the monomers (Supplementary Fig. 5B, S5C: likelihood ratio > 100).

We found that AP-1 monomers with substantive binding before KLA treatment (ATF3, Jun, and JunD) showed changes in their preference (as measured by the likelihood ratio for each motif when comparing the KLA and Vehicle TBA models) for motifs bound by upregulated TFs such as Rel, Irf3/7/8/9, Irf2 and Nfat (Fig. 5B, likelihood ratio > 10e4). Conversely, down regulated TFs were found to have reduced significance for all AP-1 monomers after 1-hour KLA treatment including Usf (Fig. 5B, likelihood ratio < 10e-4). AP-1 monomers activated after 1-hour KLA treatment (Fos, FosL2 and JunB) (Fig. 2A, 5A) also showed an affinity for the Rel, Nfat, Irf3/7/8/9 and NFκB motifs (Fig. 5B).

To assess the extent to which individual TF motifs could explain the change in binding after KLA treatment, we calculated the correlation of each motif’s score to the change in binding after KLA treatment at all loci (Fig. 5C). We found that motifs with large changes in significance when comparing the Vehicle and KLA TBA models for each monomer showed higher correlations to the change in binding after KLA treatment and that these motifs corresponded to well established TLR4 activated TFs such as Rel, NFAT, and NFκB (Fig. 5B, 5C)^11,31^. To demonstrate that combinations of TFs can better explain the change in AP-1 binding after KLA treatment, we used TBA to predict the change in binding after KLA treatment. We calculated a predicted change in binding by taking the difference of the predicted binding strength given by the Vehicle and KLA model for each monomer (Fig. 5D-F). We found that TBA could predict the change in binding after KLA treatment better than any individual motif (Fig. 5C).

### Leveraging natural genetic variation between mouse strains to validate TBA results

To validate the results of our machine learning model genome wide, we used natural genetic variation found between C57BL6/J and BALBc/J mice, which differ genetically by ~5 million single nucleotide polymorphisms (SNPs) and insertions/deletions (InDels)^42^. We have previously shown that mutations which occur within DNA binding motifs can be used to predict genetic interactions between TFs^10,34^. We performed ChIP-seq targeting expressed AP-1 monomers, ATF3, Fos, FosL2, Jun, JunB and JunD in TGEMs isolated from BALB/cJ mice. Mutations can be found in ~17% of each monomer’s binding sites, and one third of those loci show strain specific binding (fold change >2), as shown for ATF3 (Fig. 6A). These binding differences cannot be attributed to differences in the mRNA expression levels, which are highly similar (Supplementary Fig. 6A). We observed that TBA models trained on either strain could be used to predict binding in the other with no loss of predictive ability (Supplementary Fig. 6B), suggesting that each monomer, which has identical protein sequence in both strains, interacts with the same repertoire of collaborating TFs in both strains.

**Figure 6.**
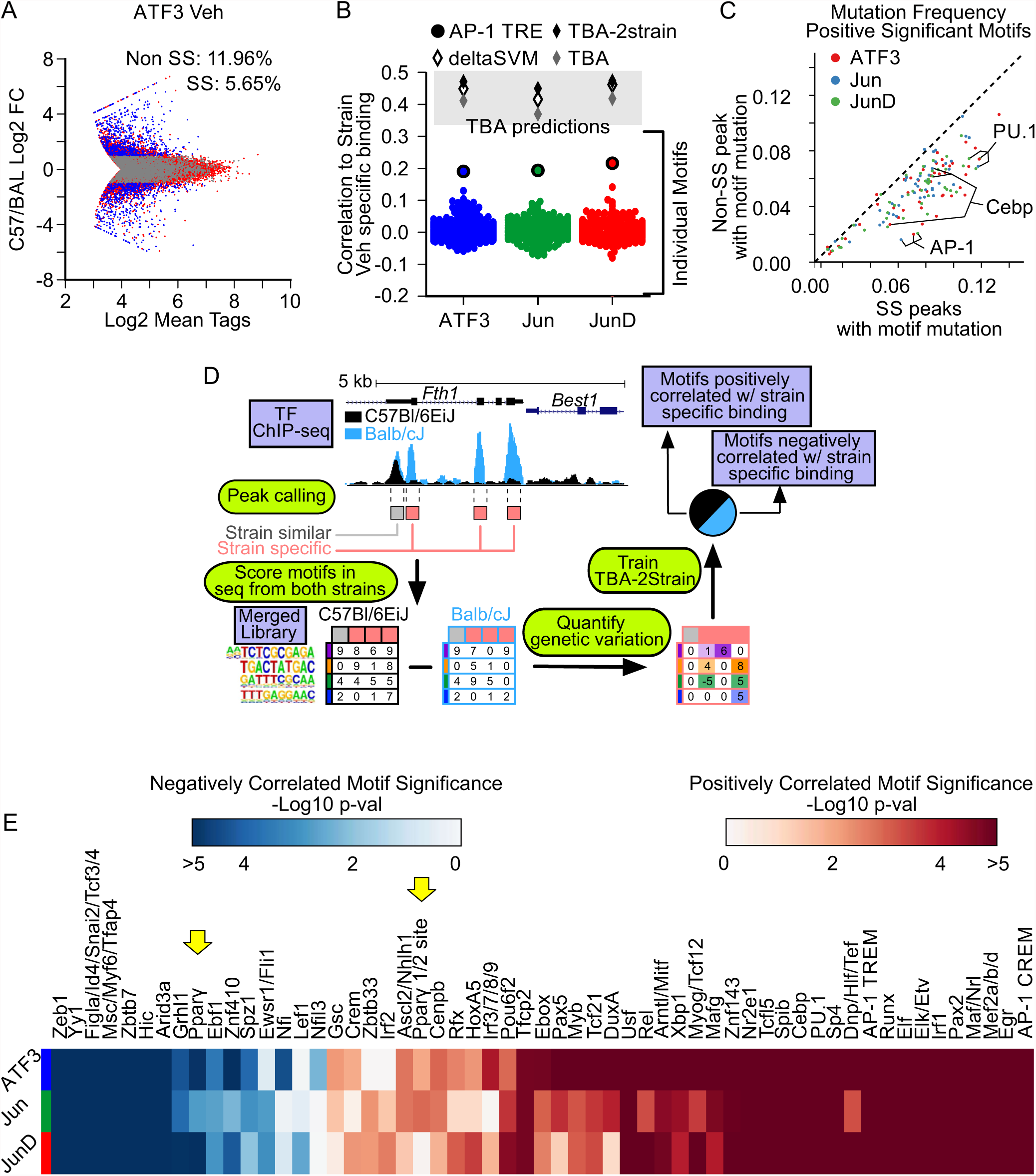
Leveraging the effects of genetic variation to validate TBA predictions in resting macrophages. **A.** Comparison of the mean strength of binding (number of quantile normalized ChIP-seq tags) for ATF3 in resting TGEMs isolated from C57Bl/6J and Balb/cJ versus the extent of strain specific binding. Loci with a mutation are indicated in blue (fold change ≥2) when there is strain specific binding and grey otherwise. **B.** Comparison of different models for predicting strain specific binding of each monomer as measured by the Pearson correlation of a model’s predictions versus the extent of strain specific binding in resting TGEMs. Models that integrate multiple motifs – deltaSVM, TBA, TBA-2Strain, are represented as diamonds. Individual motifs are indicated using round points. **C.** Frequency of mutations in significant motifs (from TBA model, p<10e-2.5) at strain specific (fold change ≥2) versus non-strain specific peaks resting TGEMs. **D.** Schematic of TBA-2Strain model. Binding sites for a transcription factor with at least one SNP or indel (red boxes) and binding sites with no mutation (grey) are identified. Next, genetic variation is quantified as the difference in the motif scores between the sequences from the two strains and then used as input to train the TBA-2Strain model to predict the extent of strain specific binding. Model weights from the trained model indicate whether a mutation in a motif is correlated with strain specific binding. **E.** Heatmap of significance values for motifs that intersected between the TBA and TBA-2Strain model for each monomer in resting TGEMs. Blue indicates motifs negatively correlated with binding and red indicates positively correlated motifs.

To assess the extent to which SNPs/InDels in individual motifs explain strain-specific binding, we calculated the difference between the best matching motif score at every loci between the strains and then calculated the Pearson Correlation to the change in binding (Fig. 6B, S6C). Mutations in individual motifs showed a weak correlation to strain specific binding (Fig. 6B, S6C). We found that motifs identified with TBA (p<10e-2.5) are enriched at strain specific peaks in comparison to non-strain specific peaks, but that mutations in any individual motif do not occur frequently enough to explain the majority of strain specific binding (Fig. 6C, S6D). We integrated the contributions of multiple motifs to strain specific binding, by weighting the motif score difference with the TBA calculated weight, and were able to predict strain specific binding with a 2-fold improvement in performance in comparison to using the AP-1 motif score (Fig. 6B, S6C)

Next, we created a variant of our model, which we call TBA-2Strain, that directly learns from genetic variation (Fig. 6D). TBA-2Strain takes genetic variation as input (quantified as the change in motif scores between the two strains) and the extent of strain specific binding for each AP-1 monomer. Using TBA- 2strain, we predicted strain specific binding at all binding sites with a mutation (Fig. 6B). In comparison to TBA, TBA-2Strain has better predictive performance (Fig 6B). This may be attributed to TBA-2Strain being able to observe sites that contain mutations but do not exhibit strain specific binding. The ability of TBA-2Strain to predict strain specific binding improves upon deltaSVM, a state of the art tool for predicting the effect the genetic variation^40^ (Fig. 6B, S6C).

We then extracted significant motifs from TBA-2Strain using the F-test (p < 0.05) and intersected these motifs with motifs identified by TBA model (Fig. 6D, 4D). We found that the motifs from both models overlapped substantially (Fig. 6D, p < 0.05, Fisher’s exact test), reinforcing the notion that dozens of motifs contribute to coordinating the targeting of AP-1 monomers. Significance values for motifs identified by both models are shown from resting and activated TGEMs(Fig. 6H, S6E). Notably, the PPARγ half-site was detected by both the TBA and TBA-2Strain models.

### Validation of PPARγ as a preferential modifier of Jun binding

TBA and TBA-2Strain predicted that PPARγ is a preferential collaborating TF specific to Jun in resting macrophages (Fig. 4A, Fig. 6E). To confirm this prediction, we performed ChIP-seq for ATF3, Jun, JunD and PPARγ in wild type and PPARγ knockout mouse TGEMs (Fig. 7 A-C)^43^. Representative browser tracks are shown for Jun binding in wild-type and PPARγ knockout macrophages (Fig. 7D). The protein expression of ATF3, Jun and JunD are unchanged in PPARγ knockout TGEMs in comparison to wild type (Fig. 7E). ChIP-seq experiments in PPARγ knockout TGEMs show a marked reduction in Jun binding (Fig. 7A). In contrast, ATF3 and JunD show little change in binding (Fig. 7B, C). We found that PPARγ bound loci where Jun binding is lost in the PPARγ knockout tended to score higher for the PPARγ half site motif in comparison to Jun bound loci that did not overlap with PPARγ binding (independent T-test p<5e-05). Collectively, these results confirm that PPARγ specifically affects Jun recruitment.

**Figure 7.**
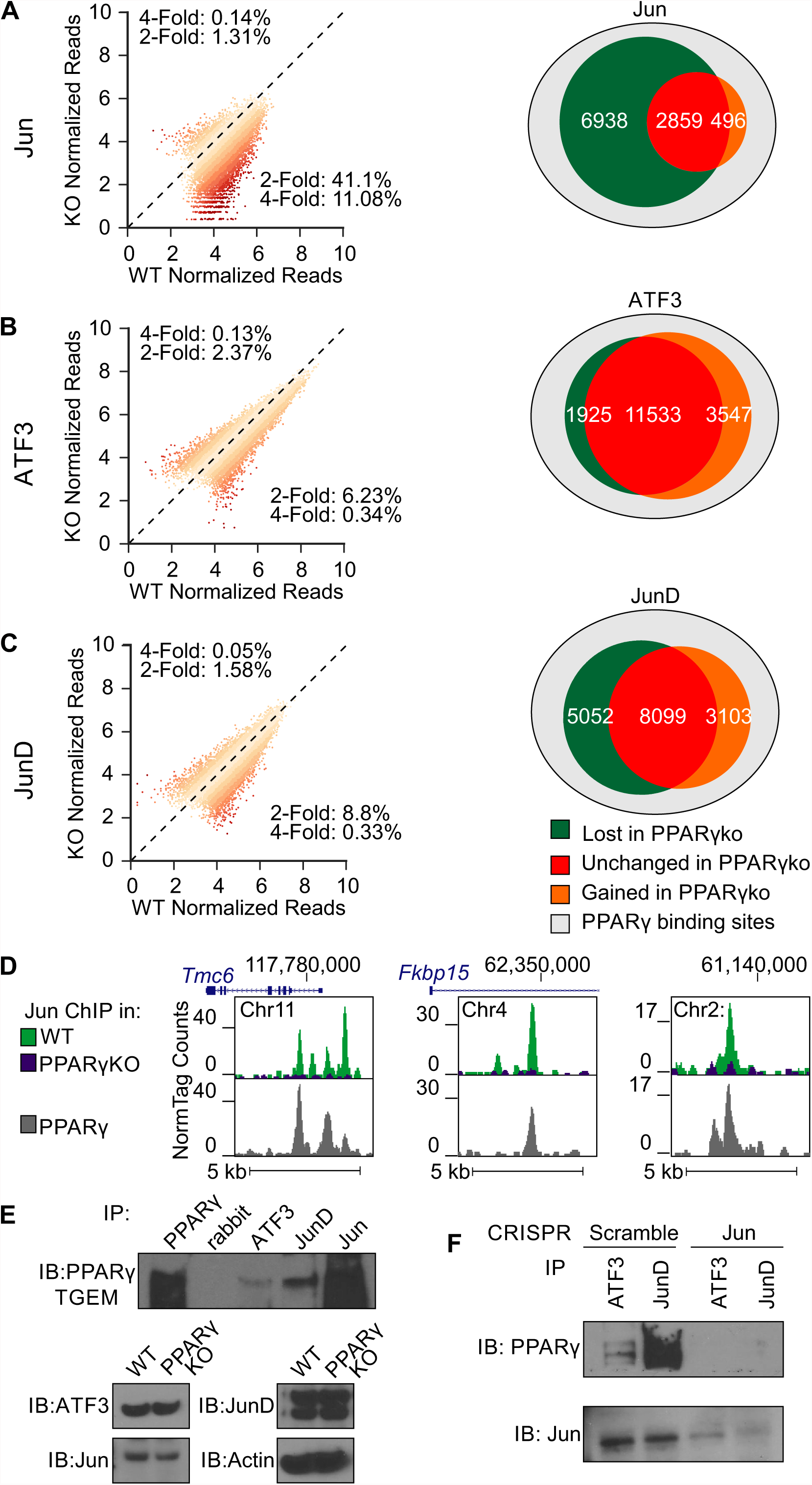
The Jun-specific DNA binding program is preferentially altered in PPARγ knockout macrophages. **A-C**. Changes in binding strength all binding sites in wild type macrophages in PPARγ-KO macrophages (left) and venn diagrams summarizing the change in binding at binding sites that overlap with PPARγ (right) for Jun **(A)**, ATF3 **(B)** and JunD **(C)**. D. Representative browser shots of Jun in WT and PPARγ-KO TGEMs and PPARγ in WT TGEMs. **E.** Western blot analysis of co-immunoprecipiation experiments between AP-1 monomers ATF3, Jun and JunD and PPARγ in TGEMs. **F.** Western blot analysis of co-immunoprecipiation experiments between AP-1 monomers ATF3 and JunD and PPARγ in scramble iBMDM or CRISPR mediated Jun knockout iBMDM

We then probed the interactions between PPARγ and AP-1 family members by co-immunoprecipitation. ATF3, Jun, and JunD co-precipitated with PPARγ (Fig. 7E). As AP-1 binds as a dimer, ATF3 and JunD may be interacting with PPARγ indirectly by dimerizing with Jun. To confirm that Jun is required for interaction of ATF3 and JunD with PPARγ, we performed Co-IP from iBMDM cells in which Jun was knocked out using CRISPR/Cas9 (Supplemental Fig. 1B). We found a loss of interaction between PPARγ and ATF3 or JunD in JunKO cells as compared to scramble control (Fig. 7F). This suggests that ATF3 and JunD do not interact with PPARγ in the absence of Jun.

## Discussion

We demonstrate that AP-1 monomers have both distinct and overlapping transcriptional functions and genome-wide binding patterns in macrophages. Monomer-specific differences in DNA binding are not due to differences in the DBD contact residues as demonstrated by ATF3 chimeras with Jun or Fos DBDs. These observations led us to hypothesize that monomer-specific DNA binding patterns result from locus-specific interactions with different ensembles of collaborating TFs. To address this question, we developed a machine learning model that identified combinations of motifs that are correlated with the binding of a TF. Through this approach, we inferred TF cooperation via the presence of DNA motifs correlated with the binding of each AP-1 monomer. Leveraging the natural genetic variation found between C57BL/6J and BALB/cJ, we confirmed that mutations in motifs predicted by TBA affect AP-1 binding. Finally, we confirmed that PPARγ plays a preferential role in coordinating Jun binding in TGEMs.

In designing our machine learning model, we optimized for interpretability. We leveraged logistic regression, a relatively simple method, to accurately predict TF binding, and we were able to extract TF motifs underlying these predictions, allowing for the generation of biological hypotheses that can be experimentally validated. A secondary benefit of this approach is that the software can be readily used without specialized computing equipment or a high level of computational understanding. To improve the ability of TBA to robustly identify motifs of interest, we programmatically curated a library that ”captures” the core of each motif, thereby mitigating collinearity, which can cause machine learning models to produce inaccurate results. By jointly weighing this library of motifs, TBA enables the detection of combinations of TF binding sites that can predict the distinct and overlapping DNA binding of families of TFs that recognize similar sequences. More broadly, TBA can be applied to predict of the effects of mutations on TF binding, and identify determinants of enhancer activation and open chromatin.

There are additional complexities in TF binding and enhancer activation we have not explored. Transcriptional regulation may be encoded by the spacing between motifs as well as the specific arrangement of motifs. Although more complex machine learning techniques can be applied to predict TF binding and chromatin state^44-46^, it is challenging to extract insights from these models. Recent neural network architectures, such as CapsuleNets, could allow modelling of these complex properties^47-49^.

Collectively, our findings suggest two classes of collaborative TFs: 1) highly ranked TFs that are strongly correlated with the binding of all AP-1 monomers, including TFs important to macrophage identity such as such as PU.1 and C/EBPs^10,11,13,50-52^ (Fig. 4A, black and grey boxes), and 2) moderately ranked TFs that specify the binding of individual AP-1 monomers (Fig. 4D, red and blue boxes). The former likely consists of TFs that play a role in opening chromatin while the latter class of TFs may allow for tuning the optimal level of transcriptional activation or response. These two classes of motifs were also seen in TLR4 activated macrophages where highly ranked motifs, such as NFκB, were correlated with the binding of all AP-1 family members (Supp Table 1), while a large set of moderately ranked motifs distinguished each AP-1 monomer (Supplementary Fig. 5C). Overall, these studies provide evidence that collaborative interactions of TFs allow a single DNA motif to be used in a wide variety of contexts, which may be a general principle for how transcriptional specificity is encoded by the genome.

## Methods

### Statistical Analyses

In Fig. 1C, differences in gene expression was tested using the independent T-test (degree of freedom=1, two-tailed) on two replicate experiments (n=2). Differentially expressed genes in Fig. 1B were identified using EdgeR^53^ with default parameters, and using the cut offs FDR¡0.05 and log2 fold change ≥2. In Fig. 2C, differences between each group (Veh, Shared, and KLA 1h) were examined using independent T-test (degree of freedom=1, two-tailed); the number of loci in each group for each monomer are as follows – ATF3 (Veh=1447, Shared=7460, KLA=6997), Jun (2390, 3751, 3401), JunD (1351, 5976, 6422). Significance for motifs in Fig. 4A was calculated using the likelihood ratio test (degree of freedom=1) comparing the predictions made by the full TBA model and the perturbed TBA model at all loci bound in Veh treated macrophages for Atf3 (n=23160), Jun (n=15548), and JunD (n=19653). Significance for motifs in Supplementary Fig. 4C was calculated using the likelihood ratio test (degree of freedom=1) comparing the predictions made by the full TBA model and the perturbed TBA model at all loci bound by JunD in GM12878 (n=7451), H1-hESC (n=12931), HepG2 (n=41318), K562 (n=47477), and SK-N-SH (38960). Significance for motifs in Fig. 5B, Supplementary Fig. 5B, and Supplementary Fig. 5C were calculated using the likelihood ratio test (degree of freedom=1) comparing the predictions made by the full TBA model and the perturbed TBA model at all loci bound in KLA treated macrophages for Atf3 (n=36745), Jun (n=17481), JunD (n=31641), Fos (n=24365), Fosl2 (n=10619), and JunB (n=13376). Significance values for Fig. 6F and S6E were calculated using the F-test; the number of loci analyzed for monomers in Vehicle treated macrophages are: ATF3 (n=4163), Jun (n=3004), and JunD (n=4148); the number of loci analyzed for monomers in KLA treated macrophages are: Atf3 (n=4577), Jun (n=3232), JunD (n=4366), Fos (n=4477), and JunB (n=3616).

### Generating Custom Genome for BALB/cJ

A custom genome for BALB/cJ by replacing invariant positions of the mm10 genome with alleles reported by the Mouse Genomes Project (version 3 VCF file)^42^. For C57BL/6J the mm10 reference genome from the UCSC genome browser was used. To allow for comparisons between BALB/cJ and C57BL/6J during analysis, the coordinates for the custom genome for BALB/cJ was shifted to match the positions of the mm10 reference genome using MARGE^34^. We did not analyze any reads that fell within deletions in BALB/cJ. Reads that overlapped with an insertion were assigned to the last overlapping position in the reference strain.

### Analysis of ChIP-seq Peaks

Sequencing reads from ChIP-seq experiments were mapped to the mm10 assembly of the mouse reference genome (or the BALBc/J custom genome) using the latest version of Bowtie2 with default parameters^54^. Mapped ChIP-seq reads to identify putative transcription factor binding sites with HOMER^55^ findPeaks command (with parameters -size 200 -L 0 -C 0 -fdr 0.9), using the input ChIP experiment corresponding to the treatment condition. In order to reduce the number of false positive peaks, we calculated the Irreproducible Discovery Rate (IDR) at each peak (using version 2.0.3 of the idr program) with the HOMER peak score calculated for each replicate experiment as the input to IDR and then filtered all peaks that had IDR ≥ 0.05^56^. De novo motifs were calculated with the HOMER findMotifsGenome.pl command with default parameters. Enrichment of de novo motifs was calculated using the findKnownMotifs.pl program in HOMER with default parameters.

Quantification of RNA Expression Reads generated from RNA-seq experiments were aligned to the mm10 mouse reference genome (or the BALBc/J custom genome) using STAR aligner with default parameters^57^. To quantify the expression level of each gene, we calculated the Reads Per Kilobase of transcript per Million mapped reads (RPKM) with the reads that were within an exon. Un-normalized sequencing reads were used identify differentially expressed genes with EdgeR^53^; we considered genes with FDR < 0.05 and a change in expression between two experimental conditions two fold or greater differentially expressed. To quantify the expression of nascent RNAs we annotated our ChIP-seq peaks with the number of GRO-seq reads (normalized to 10 million) that were within 500 bps of the peak center using the HOMER annotatePeaks.pl command.

### TBA Model Training

For each AP-1 monomer under each treatment condition, we trained a model to distinguish binding sites for each monomer from a set of randomly selected genomic loci. The set of random background loci used to train each model was selected according to the following criteria: 1) the GC content distribution of the background loci matches the GC content of the binding sites for a given monomer, 2) contain no ambiguous or unmappable positions, and 3) the number of background sequences matches the number of binding sites k. For each of the sequences in the combined set of the binding sites and background loci, we calculated the highest log-odds score (also referred to as motif score) for each of the *n* motifs that will be included in the model^58^ Motif matches in both orientations were considered. Log-odds scores less than 0 were set to 0. Per standard preprocessing procedures prior to training a linear model, we standardized the log-odds scores for each motif, scaling the set of scores for each motif so that the mean value is 0, and the variance is 1. Standardization scales the scores for all motifs to the same range (longer motifs have a larger maximum score) and also helps to reduce the effect of multi-collinearity on the model training. And so, the features used for training our model is an *n* by 2*k* matrix of log-odds scores standardized across each row. To generate the corresponding array of labels, we assigned each binding site a label of 1 and each background loci a label of 0. Using this feature matrix, and label array, we trained weights for each motif using an L1 penalized logistic regression model as implemented by the scikit-learn Python package^59^. Motif weights shown in our analysis are the mean values across five rounds of cross validation, using 80% of the data for training and 20% for testing in each round. Models were trained for ChIP-seqs generated in this study as well as data downloaded from the NCBI Gene Expression Omnibus (accession number GSE46494) and the ENCODE data portal (www.encodeproject.org).

### Quantification of Multiple Collinearity

To assess the extent multi-collinearity in the motif score features we used train our models, we took each feature matrix corresponding to each experiment and calculated the Variance Inflation Factor (VIF) for each motif^38^. To calculate the VIF, we first determine the coefficient of determination, R2, for each motif by regressing the log-odds scores for one motif against the log-odds scores of the remaining motifs. Next using the coefficient of determination, the tolerance for each motif can be calculate as the difference between 1 and the coefficient of determination (1−*R*^2^). The VIF is the reciprocal of the tolerance 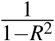. We used the linear model module of sklearn Python package to calculate the coefficient of determination.

### Motif Clustering and Merging

We scored the similarity of all pairs of DNA sequence motifs by calculating the Pearson correlation of the aligned position probability matrices (PPMs) corresponding to a given pair of motifs. PPMs were first aligned using the Smith-Waterman alignment algorithm^60^. Shorter motifs are padded with background frequency values prior to alignment. Gaps in the alignment were not allowed and each position in the alignment was scored with the Pearson correlation. The Pearson Correlation was then calculated using the optimal alignment. Next, sets of motifs that have PPMs with a Pearson correlation of 0.9 or greater were merged by iteratively aligning each PPM within the set, and then averaging the nucleotide frequencies at each position.

### Assessing Significance of Motifs for TBA

p-values for TBA were calculated using the log likelihood ratio test. Each motif was removed from the set of features used to train a perturbed TBA model (using five-fold cross validation). We then used the full model (containing all motifs) and the perturbed model to calculate the likelihood of observing binding on all binding sites and background sequences for a given monomer and all the background regions. The difference in the likelihoods calculated by the full model and the perturbed model was then used to perform the chi-squared test for each motif. The chi-squared test was performed using the scipy python package^61^

### Comparison to other Methods

BaMM motif and gkm-SVM were both run with default parameters. We used the latest version of the large scale gkm-SVM, LS-GKM, and BaMM motif^39,62^. Both models were trained using five-fold cross validation. Model performance was scored using roc auc score and precision score functions from the metrics module of sklearn.

### Predicting changes in AP-1 binding after one-hour KLA treatment

To predict the change in binding after KLA treatment, we leveraged the motif weights learned for each of the *n* motifs (*w_n_*) by a TBA model trained on the Vehicle treated data (*W_v_eh* = [*w_veh,_*_1_,…*w_veh,n_*]) and a TBA model trained on the one-hour KLA treated data (*W_k_la* = [*w_kla,_*_1_,…*w_kla,n_*]) for each AP-1 monomer. The predicted change in binding for each sequence is then then the difference between the dot product of the standardized motif scores calculated for the sequence each of the *k* binding sites (*S_k_* = [*s*_1*,k*_,…, *s_n,k_*]) with the KLA motif weights and the dot product of the motif scores and the Veh motif weights (Δ*_kla_*_−*veh,k*_ = *W_kla_* ·*S_k_* − *W_veh_* · *S_k_*). Predictions were made for all genomic loci that intersected with a peak for one of the AP-1 monomers in either the vehicle or KLA treatment condition.

### Predicting strain specific binding with TBA

To predict strain specific binding, we leveraged the motif weights learned for each of the *n* motifs (*w_n_*) by a TBA model (*W* = [*w*_1_,…,*w_n_*]) for each AP-1 monomer using the C57BL/6J data, and the motif scores calculated for each of the *k* binding sites using the genomic sequence for C57BL/6J and BALBc/J (*S_C_*_57*,k*_ = [*s_C_*_57,1*,k*_,…, *s_C_*_57*,n,k*_]*, S_BAL,k_* = [*s_BAL,_*_1*,k*_,…, *s_BAL,n,k_*]). Next, we computed the difference of the motif scores for C57BL6/J and BALBc/J ([*D_n_* = [*s_C_*_57*,n,*1_ − *s_BAL,n,_*_1_,…, *s_C_*_57*,n,k*_ − *s_BAL,n,k_*]) and then standardized the score differences for each motif across all the *k* binding sites that had a mutation when comparing BALBc/J to C57BL/6J, yielding standardized motif score differences for each binding site (*Z_n_* = *standardize*(*D_n_*) = [*z_n,_*_1_,…,*z_n,k_*]). Finally, we then made a prediction for strain specific binding by computing the dot product of the motif weights and the standardized difference of the motif scores between C57BL6/EiJ and BALBc/J for the k^th^ mutated binding site (Δ*_C_*_57−*BAL*_ = *W* · [*z*_1*,k*_,…, *z_n,k_*]).

### TBA-2Strain Model Training

For each genomic loci that intersected with a peak for one of the AP-1 monomers, in either C57BL/6J or BALBc/J, we calculated the highest log-odds score for each of the *n* motifs that will be included in the model, using the genomic sequence from both strains, yielding a two sets of motif scores for each of the *k* binding sites (*S_C_*_57*,k*_ = [*s_C_*_57,1*,k*_,…, *s_C_*_57*,n,k*_]*, S_BAL,k_* = [*s_BAL,_*_1*,k*_,…, *s_BAL,n,k_*]). Motif matches in both orientations were considered. Log-odds scores less than 0 were set to 0. Using the motif scores, we compute the standardized difference of the motif scores across the two strains as described in the above section (*Z_n_* = [*z_n,_*_1_,…*, z_n,k_*]).And so, the features used for training our model is an *n* by *k* matrix of log-odds scores standardized across each row. Next, we calculated the log2 fold ratio of the number of ChIP-seq reads in C57BL/6J compared to BALBc/J to represent the extent of strain specific binding. Using this feature matrix, and setting the log2 fold ratio of binding between the two strains as the dependent variable, we trained weights for each motif using linear regression as implemented by the scikit-learn Python package. Motif weights shown in our analysis are the mean values across five rounds of cross validation, using 80% of the data for training and 20% for testing in each round. Predictions for strain specific binding can be made using the calculated weights following the procedure in the previous section.

### Code Availability

All algorithms relating to training and testing our model, TBA, has been implemented using Python. Source code and executable files are available at: https://github.com/jenhantao/tba.

### ChIP protocol

Protein A and G Dynabeads 50/50 mix from Invitrogen are sued for ChIP (10001D, 10003D). IP mix consists of 20ul beads/2ug antibody per 2 million cell ChIP. For preparation, beads were washed with 0.5%BSA-PBS (BSA:, then incubate beads-antibody with 0.5%BSA-PBS for at least 1h on rotator (4oC). Wash 2X with 0.5%BSA-PBS, then resuspend in dilution buffer. Double Crosslinking for ChIP. Media was decanted from cells in 10 cm plates, wash once briefly with PBS (RT). Disuccinimidyl glutarate (Pierce Cat # 20593) (diluted in DMSO at 200mM)/PBS (RT) was used for 10 min. Then Formaldehyde was added to a final concentration of 1% for an additional 10 min. Reaction was quenched with 1:10 Tris pH 7.4 on ice. Cells were collected and washed twice with cold PBS, spinning at 1000 rcf for 5 min. Nuclei Isolation and Sonication. Resuspend cell pellets in 1 ml of Nuclei Isolation Buffer (50 mM Tris-Ph 8.0, 60 mM KCl, 0.5% NP40) + PI and incubate on ice for 10 minutes. Centrifuge 2,000 g for 3 minutes at 4° C. Resuspend nuclei in 200 ul of fresh Lysis Buffer (0.5% SDS, 10 mM EDTA, 0.5 mM EGTA, 50 mM Tris-HCl (ph8))+ PI. Sonication. Nuclei were then sonicated (10 million cells) for 25 minutes in a Biorupter (settings=30 seconds=On, 30 seconds=Off, Medium) using thin wall tubes (Diagenode Cat# C30010010). After sonication spin max speed for 10 minute at 4C. ChIP set up. Sonicated DNA was diluted 5X with 800 Dilution Buffer (1% Triton, 2mM EDTA, 150 mM NaCl, 20 mM Tris-HCl (ph8), 1X Protease Inhibitors). An aliquot is removed for input samples (5%). Samples ON at 4° C while rotating. Washing. ChIP are washed 1X with TSE I (20mM Tris-HCl pH7.4, 150mM NaCl, 0.1%SDS, 1% Triton X-100, 2mM EDTA), 2X with TSE III (10mM Tris-HCl pH7.4, 250mM LiCl, 1%IGEPAL, 1%Deoxycholate, 1mM EDTA), 1X with TE+0.1%TritonX-100, transfer to new tube and then wash another time with TE+0.1%TritonX-100. Elution. Elute with 200 µL Elution Buffer (1% SDS, 10mM Tris pH7.5) for 20 minutes at RT, shaking on the vortex or a nutator or rotator. De-crosslinking. Add 10 µL of 5 M NaCl and incubate ON at 65° C (or at least 8 hours). Clean up samples using Zymo ChIP DNA Clean and Concentrator. Elute in 100µL. Take 40µL and proceed to library prep protocol.

### PolyA RNA Isolation and Fragmentation

RNA isolation. RNA was isolated using TRIZOL-reagent (ambion cat# 15596018) and DIRECT-ZOL RNA mini-prep kit (cat# 11-330MB). Poly-A RNA isolation. Use 0.2 Total RNA as starting material for ideal mapping efficiency and minimal clonality. Collect 10 µL oligo (dT) (NEB cat# S1419S) beads per RNA sample. Beads were washed twice with 1x DTBB (20mM Tris-HCl pH7.5, 1M LiCl, 2mM EDTA, 1% LDS, 0.1% Triton X-100). Beads were resuspended in 50µL of 2x DTBB. 50µL of beads were mixed with 50µL RNA and Heated to 65°C for 2 min. RNA-beads were then incubated for 10 min at RT while rotating. RNA-beads were then collected on a magnet and washed 1x each with RNA WB1 (10mM Tris-HCl pH7.5, 0.12 M LiCl, 1mM EDTA, 0.1% LDS, 0.1% Triton X-100)and WB3 (10mM Tris-HCl pH7.5, 0.5M LiCl, 1mM EDTA). Add 50µL Tris-HCl pH7.5 and heat to 80°C for 2 min to elute. Collect RNA and perform a second Oligo-dT bead collection. After washing the second collection, instead of eluting was 1X with 1X First strand buffer (250 mM Tris-HCl (pH 8.3), 375 mM KCl, 15 mM MgCl2 (ThermoFisher SSIII kit Cat# 18080093). Fragmentation. Then Add 10µL of 2X First strand buffer plus 10mM DTT and fragment DNA at 94°C for 9 min. Collect beads on magnet and transfer eluate containing fragmented mRNA to a new PCR strip. Should recover 10 µL fragmented RNA. First strand synthesis. We mixed fragmented RNA with 0.5µL Random Primer (3 µg/µL) Life Tech #48190-011, 0.5µL oligo-dT (50uM from SSIII kit), 1µL dNTPs (10mM Life Tech, cat 18427088) and 0.5µL SUPERase-In (ThermoFisher Cat#AM2696) and heat 50°C for 1 min. Immediately place on ice. We then added 5.8µL ddH2O, 0.1 µL Actinomycin (2ug/µL Sigma cat#A1410), 1µL DTT (100mM Life Tech cat# P2325), 0.2µL of 1% Tween and 0.5 µL of Superscript III and incubate 25°C for 10 min, then 50°C for 50 min. Bead clean up. We added 36 µL of RNAClean XP (ampure XP) and mixed, incubating for 15 min on ice. The beads were then collected on a magnet and washed 2X with 75% ethanol. Beads were then air-dried for 10 min and elute with 10 µL nuclease free H2O. Second strand synthesis. 10µL of cDNA/RNA was mixed with 1.5µL 10X Blue Buffer (Enzymatics cat# B0110L), 1µL dUTP/dNTP mix (10mM Affymatrix cat# 77330), 0.1µL dUTP (100mM Affymatrix cat# 77206), 0.2µL RNase H (5U/µL Enzymatics cat# Y9220L), 1µL DNA polymerase I (10U/µL Enzymatics cat#P7050L), 0.15 µL 1% Tween-20 and 1.05µL nuclease free water. Reaction was incubated at 16°C for 2.5 hours. Bead clean up. DNA was purified by adding 1µL Seradyn “3 EDAC” SpeedBeads (Thermo 6515-2105-050250) per reaction in 28µL 20% PEG8000/2.5 M NaCl (13% final concentration) and incubating at RT for 10min. Beads were then collected on a magnet and washed 2X with 80% Ethanol. Beads were air-dried for 10min and eluted in 40µL of nuclease free water. DNA is ready for library prep.

### Library Prep Protocol

dsDNA End Repair. We mixed 40µL of DNA from ChIP or RNA protocols with 2.9µL of H2O, 0.5µL 1% Tween-20, 5µL 10X T4 ligase buffer (Enzymatics cat# L6030-HC-L), 1µL dNTP mix (10 mM Affymetrix 77119), 0.3 µL T4 DNA pol (Enzymatics P7080L), 0.3µL T4 PNK (Enzymatics Y9040L), 0.06µL Klenow (Enzymatics P7060L) and incubated for 30min at 20°C. 1µL of Seradyn “3 EDAC” SpeedBeads (Thermo 6515-2105-050250) in 93 µL 20% PEG8000/2.5 M NaCl (13% final) was added and incubated for 10 min. Bead clean-up. Beads were collected on a magnet and washed 2X with 80% ethanol. Beads were air-dried for 10 min and then eluted in 15µL ddH2O. dA-Tailing. DNA was mixed with 10.8µL ddH2O, 0.3µL 1% Tween-20, 3µL Blue Buffer (Enzymatics cat# B0110L), 0.6µL dATP (10mM Tech 10216-018), 0.3µL Klenow 3’- 5’ Exo (Enzymatics P7010-LC-L) and incubated for 30min at 37°C. 55.8µL 20% PEG8000/2.5 M NaCl (13% final) was added an incubated for 10 min. Then bead clean up was done. Beads were eluted in 14µL. Y-Shape Adapter Ligation. Sample was mixed with 0.5µL of a BIOO barcode adapter (BIOO Scientific cat# 514104), 15µL Rapid Ligation Buffer (Enzymatics cat@ L603-LC-L), 0.33µL 1% Tween-20 and 0.5µL T4 DNA ligase HC (Enzymatics L6030-HC-L) and incubated for 15 min at RT. 7 µL of 20% PEG8000/2.5 M NaCl was added and incubated for 10min at RT. Bead clean up was performed and beads were eluted in 21µL. 10µL was then used for PCR amplification (14 cycles) with IGA and IGB primers (AATGATACGGCGACCACCGA, CAAGCAGAAGACGGCATACGA).

### GRO-seq

Nascent transcription was captured by global nuclear run-on sequencing (GRO-seq). Nuclei isolation. Nuclei were isolated from TGEMs using hypotonic lysis (10 mM Tris-HCl (pH 7.5), 2 mM MgCl2, 3 mM CaCl2; 0.1% IGEPAL CA-630) and flash frozen in GRO-freezing buffer (50 mM Tris-HCl (pH 7.8), 5 mM MgCl2, 40% Glycerol). Run-on. 3-5 × 106 BMDM nuclei were run-on with BrUTP-labelled NTPs with 3x NRO buffer (15mM Tris-Cl (pH 8.0), 7.5 mM MgCl2, 1.5 mM DTT, 450 mM KCl, 0.3 U/µL of SUPERase In, 1.5% Sarkosyl, 366 uM ATP, GTP (Roche), Br-UTP (Sigma 40 Aldrich) and 1.2 uM CTP (Roche, to limit run-on length to ~40 nt)). Reactions were stopped after five minutes by addition of 500 µL Trizol LS reagent (Invitrogen), vortexed for 5 minutes and RNA extracted and precipitated as described by the manufacturer. Fragmentation. RNA pellets were resuspended in 18 ul ddH2O + 0.05% Tween (dH2O+T) and 2 ul fragmentation mix (100 mM ZnCl2, 10 mM Tris-HCl (pH 7.5)), then incubated at 70°C for 15 minutes. Fragmentation was stopped by addition of 2.5 ul 100 mM EDTA. BrdU enrichment. BrdU enrichment was performed using BrdU Antibody (IIB5) AC beads (Santa Cruz, sc-32323 AC, lot #A0215 and #C1716). Beads were washed once with GRO binding buffer (0.25xsaline-sodium-phosphate-EDTA buffer (SSPE), 0.05% (vol/vol) Tween, 37.5 mM NaCl, 1 mM EDTA) + 300 mM NaCl followed by three washes in GRO binding buffer and resuspend as 25% (vol/vol) slurry with 0.1 U/µL SUPERase-in. To fragmented RNA, 500 µL cold GRO binding buffer and 40 µL equilibrated BrdU antibody beads were added and samples slowly rotated at 4°C for 80 minutes. Beads were subsequently spun down at 1000xG for 15 seconds, supernatant removed and the beads transferred to a Millipore Ultrafree MC column (UFC30HVNB; Millipore) in 2x 200 µL GRO binding buffer. The IP reaction was washed twice with 400 µL GRO binding buffer before RNA was eluted by incubation in 200 µL Trizol LS (Thermo Fisher) under gentle agitation for 3 minutes. The elution was repeated a second time, 120 µL of dH2O+T added to increase the supernatant and extracted as described by the manufacturer. End repair and decapping. For end-repair and decapping, RNA pellets were dissolved in 8 ul TET (10 mM Tris-HCl (pH 7.5), 1 mM EDTA, 0.05 % Tween20) by vigorous vortexing, heated to 70°C for 2 minutes and placed on ice. After a quick spin, 22 ul Repair MM (3 ul 10x PNK buffer, 15.5 ul dH2O+T, 0.5 ul SUPERase-In RNase Inhibitor (10 U), 2 ul PNK (20U), 1 ul RppH (5U)) was added, mixed and incubated at 37°C for 1 hour. 5’ Phosphorylation. To phosphorylate the 5’end, 0.5 ul 100 mM ATP was subsequently added and the reactions were incubated for another 45 minutes at 37°C (the high ATP concentration quenches RppH activity). Following end repair, 2.5 ul 50 mM EDTA was added, reactions mixed and then heated to 70°C for 2 minutes before being placed on ice. A second BrdU enrichment was performed as detailed above. Adapter Ligation. RNA pellets were dissolved in 2.75 TET + 0.25 Illumina TruSeq 3’Adapter (10 M), heated to 70°C for 2 minutes and placed on ice. 7 of 3’MM (4.75 50% PEG8000, 1 10x T4 RNA ligase buffer, 0.25 SUPERase-In, 1 T4 RNA Ligase 2 truncated (200U; NEB)) was added, mixed well and reactions incubated at 20°C for 1 hour. Reactions were diluted by addition of 10 TET + 2 ul 50 mM EDTA, heated to 70°C for 2 minutes, placed on ice and a third round of BrdUTP enrichment was performed. RNA pellets were transferred to PCR strips during the 75% ethanol wash and dried. Samples were dissolved in 4 TET (10 mM Tris-HCl (pH 7.5), 0.1 mM EDTA, 0.05% Tween 20) + 1 10 M reverse transcription (RT) primer. To anneal the RTprimer, the mixture was incubated at 75°C for 5 minutes, 37°C for 15 minutes and 25°C for 10 minutes. To ligate the 5’ Illumina TruSeq adapter, 10 5’MM (1.5 ddH2O + 0.2% Tween20, 0.25 denaturated 5’TruSeq adapter (10 M), 1.5 10x T4 RNA ligase buffer, 0.25 SUPERase-In, 0.2 10 mM ATP, 5.8 50% PEG8000, 0.5 T4 RNA ligase 1 (5U; NEB)) was added and reactions were incubated at 25°C for 1 hour. Reverse Transcription. Reverse transcription was performed using Protoscript II (NEB) (4 5x NEB FirstStrand buffer (NEB; E7421AA), 0.25 SUPERase-In, 0.75 Protoscript II (150U; NEB)) at 50°C for 1 hour. After addition of 30 PCR MM (25 2X LongAmp Taq 2X Master Mix (NEB), 0.2 100 M forward primer, 2.8 5M Betaine and 2 10 M individual barcoding primer), mixtures were amplified (95°C for 3 minutes, (95°C for 60 seconds, 62°C for 30 seconds, 72°C for 15 seconds) x13, 72°C for 3 minutes). PCR reactions were cleaned up using 1.5 volumes of SpeedBeads™ (GE Healthcare) in 2.5M NaCl/20% PEG8000 Library selection and cleaning. Libraries were size selected on a PAGE/TBE gels to 160-225 base pairs. Gel slices were shredded by spinning through a 0.5 ml perforated PCR tube placed on top of a 1.5 ml tube. 150 ul Gel EB (0.1% LDS, 1M LiCl, 10 mM Tris-HCl (pH 7.8)) was added and the slurry incubate under agitation overnight. To purify the eluted DNA, 700 ul Zymogen ChIP DNA binding buffer was added into the 1.5 ml tube containing the shredded gel slice and the Gel EB, mixed by pipetting and the slurry transferred to a ZymoMiniElute column. Samples were first spun at 1000xG for 3 minutes, then 10,000xG for 30 seconds. Flow through was removed, and samples washed with 200 ul Zymo WashBuffer (with EtOH). Gel remainders were removed by flicking and columns washed by addition of another 200 ul Zymo WashBuffer (with EtOH). Flow through was 42 removed, columns spun dry by centrifugation at 14,000xG for 1 minute and DNA eluted by addition of 20 ul pre-warmed Sequencing TET (10 mM Tris-HCl (pH 8.0), 0.1 mM EDTA, 0.05% Tween 20). Libraries were sequenced.

### Western Blotting

Cells were lysed with Igepal lysis buffer (50mM Tris pH8.0, 150mM NaCl, 0.5% Igepal) and protein concentrations were determined with BioRad protein assay reagent using BSA as a standard. Proteins were separated on NuPage 4-12% Bis-Tris gradient gels (Invitrogen) and transferred onto a nitrocellulose membrane (Amersham). Membranes were blocked in TBS with 0.1% Tween-20 and 5% BSA. Membranes were blotted with the indicated primary overnight at 4oC. Horseradish peroxidase conjugated secondary antibodies were detected using ECL plus western blotting detection system (Amersham).

### Animals and Cell Culture

TGEMs were collected 3 days after injection from male 8 week C57Bl/6J, or BALB/cJ mice, and plated at 20 × 106 cells per 15 cm Petri dish in DMEM plus 10% FBS and 1x penicillin-streptomycin. One day after plating, cells were supplemented with fresh media and treated with PBS (Veh) or 100 ng/ml KLA for 1 hour, and then directly used for downstream analyses. All animal experiments were performed in compliance with the ethical standards set forth by University of California, San Diego’s Institutional Annual Care and Use Committee (IUCAC).

## Data Availability

Data generated for this study has been deposited to the NCBI Gene Expression Omnibus (GEO) under the accession number GSE111856. Previously published data was downloaded from GEO (accession number GSE46494) and the ENCODE data portal (https://www.encodeproject.org).

## Acknowledgements

We thank L. Van Ael for assistance with manuscript preparation and J. Collier, M. Pasillas and Z. Ouyang for technical assistance. These studies were supported by NIH grants DK091183, CA17390 and GM085764 and Leducq Transatlantic Network grant 16CVD01 to CKG. DNA sequencing was supported by NIH grant DK063491. SHD is a CRI-Irvington Postdoctoral Fellow. TS was supported by the Swedish Society for Medical Research. GJF was supported by a Canadian Institute of Health Research Postdoctoral Fellowship, FME-135475.

## Author contributions statement

GJF, JT, and CKG conceived the study. GJF, JT, EMW, SD, JDS, TS, NJS and VL performed experiments. JT, ZS, CB and GJF analyzed data. GJF, JT and CKG wrote the manuscript with contributions from CB.

## Disclosure Declaration

The authors declare no conflict of interests.

